# Application of Down-Phase Targeted Auditory Stimulation During Sleep in a Home Setting: A Feasibility Study Across Seven Consecutive Nights

**DOI:** 10.1101/2025.03.21.644521

**Authors:** Corinne Eicher, Cristina Gallego Vázquez, Cinzia Schmid, Golo Kronenberg, Hans-Peter Landolt, Erich Seifritz, Giulia Da Poian, Reto Huber

## Abstract

**Introduction:** Sleep deprivation, also known as “wake therapy”, has long been recognized as a powerful antidepressant. Phase-targeted auditory stimulation (PTAS) has been suggested as an auspicious non-invasive nocturnal substitute for sleep deprivation. Down-PTAS with stimuli presentation during the down-phase of slow waves, in particular, may have therapeutic potential to improve mood by selectively reducing slow-wave activity (SWA). With down-PTAS being more nuanced than sleep deprivation, its effects presumably develop over multiple nights, thus necessitating transfer from sleep laboratory to home settings. Therefore, in this study, we investigated the technical feasibility, tolerability, and potential risks associated with a wearable device employed for down-PTAS in an unsupervised home setting.

**Methods:** We recorded frontal EEG using the MHSL Sleepband Version 3 (MHSL-SB) in five healthy participants (23.8 ± 0.8 years, three women) over seven consecutive nights with (STIM) and without (SHAM) tone application at their homes. Tones were delivered shortly before the negative peak of slow waves during N2/N3 sleep, using alternating 10-s ON-OFF windows. Sleep staging followed American Academy of Sleep Medicine (AASM) guidelines. Of the 67 available sleep recordings, we excluded three due to parameter adjustments and another six for technical issues, leaving 58 sleep recordings (29 SHAM, 29 STIM) for further analyses. Low SWA (0.5-2 Hz, lSWA) was computed across the entire night, ON-OFF windows, and sleep cycles. Time-frequency analyses were performed time-locked to stimulus onset. We computed linear mixed effect models with condition (STIM vs. SHAM) as a fixed effect and random participant intercepts.

**Results:** Data quality was sufficient for analyses in 87% of the available sleep recordings, with an average of over 1500 correctly delivered stimuli per recording. Down-PTAS did not affect sleep architecture, but it reduced lSWA primarily during the first sleep cycle when sleep pressure and lSWA were highest, and particularly in OFF windows. Additionally, stimulation elicited a K-complex-like auditory evoked response, aligning with previous laboratory findings.

**Conclusion:** Our results demonstrate the successful implementation of down-PTAS in a home setting, confirming its feasibility for long-term, unsupervised use. The K-complex-like auditory evoked response may mask potential reductions in lSWA during ON windows, posing a scientific analytical challenge. Taken together, future clinical research should now assess the effects of down-PTAS in depressed patients, in whom reducing lSWA may partly mimic sleep deprivation.

## Introduction

Most studies aiming to modulate sleep focus on deepening sleep to enhance restorative processes, such as improving memory and cognitive function [1–4]. In certain conditions, however, reducing sleep depth may also offer therapeutic benefits, with one promising application being the treatment of major depression [5–12].

Currently, there are no specific pharmacological approaches to this end. Total sleep deprivation [5,6,8,10] and selective slow-wave sleep (SWS) deprivation [9] have demonstrated significant antidepressant effects, but may be associated with negative consequences for cognitive function [13–17]. Suggesting a direct link between slow waves and the antidepressant response, greater symptom relief following SWS deprivation was associated with higher baseline slow-wave dissipation and a stronger rebound during recovery sleep [9]. Phase-targeted auditory stimulation (PTAS) offers a more targeted approach for modulating slow waves [1–4,18,19], potentially overcoming the limitations of total and SWS deprivation.

The modulation of slow waves through PTAS during non-rapid eye movement (NREM) sleep has been shown to influence slow-wave activity (SWA) with significant functional implications, and there is evidence that phase-targeting plays an important role. A single night of up-PTAS, where auditory stimuli are presented during the up-phase of an ongoing slow wave, compared to a control night without stimulation, selectively enhanced slow oscillation rhythm (<1 Hz) and improved declarative memory consolidation in healthy adults [4]. By contrast, down-PTAS with stimulus presentation during the down-phase of slow waves not only reduced power in this frequency range, but also failed to confer any memory benefits of sleep slow oscillations, highlighting the potential clinical significance of phase-locked modulation [4]. In line with this observation, a local reduction of SWA over the sensorimotor area, induced by continuous down-PTAS, interfered with restoration, leading to impaired motor task performance and hindered the brain’s capacity to undergo neuroplastic changes [18]. On the other hand, two recent independent studies using a windowed stimulation protocol with 6– and 10-second window length reported an increase in SWA during down-PTAS in ON windows [1,3]. In this approach, stimuli were delivered only within designated ON windows and disabled during subsequent OFF windows, rather than continuously. This windowed stimulation protocol resulted in a consistent, K-complex-like response to isolated stimuli (≥5 s interstimulus interval, ISI), regardless of the targeted phase, but caused a differential SWA response between up– and down-PTAS over longer timescales [1].

In contrast to total sleep deprivation, which produces immediate and pronounced antidepressant effects [5–12], the more specific and potentially local SWA reduction induced by down-PTAS is expected to have milder effects, necessitating prolonged stimulation that is ultimately difficult to implement in a sleep laboratory setting. Therefore, transferring down-PTAS from a controlled laboratory to a home setting is crucial. The MHSL Sleepband Version 3 (MHSL-SB) was specifically designed for this purpose and has already been used for long-term up-PTAS in a home setting [20].

Consequently, we investigated the feasibility of unsupervised recording and down-PTAS over seven consecutive nights in healthy adults using the MHSL-SB in their homes. Specifically, we evaluated the technical feasibility, the tolerability of continuous application, and the device’s ability to elicit electrophysiological responses under real-world, uncontrolled conditions.

## Methods

### Participants

Five healthy young adults (mean age: 23.8 ± 0.8 years, three women) were recruited through flyers and word of mouth and participated between November 2021 and March 2022 in this feasibility study. During a screening visit, participants were assessed for the following inclusion criteria: age between 18 and 65 years, no current or past depression, good general health, motivation to participate, ability to follow study procedures, no aversion against technology, and ability to provide informed consent.

Exclusion criteria included pregnancy or lactation, unstable medical conditions, diagnosed sleep apnea or other sleep disorders, epilepsy, inflammatory central nervous system diseases, intracranial space-occupying lesions, substance use disorders, or hearing impairments. Additionally, individuals with a history of traumatic brain injury (except for concussion), prior neurosurgical procedures, shift work, or skin allergies to adhesive dressing or tape were excluded.

All participants provided written informed consent. The study was conducted at a single centre in Zurich in accordance with the Declaration of Helsinki and was approved by the ETH Zurich Ethics Commission (EK-2021-N-153).

### Study Design

The study included two one-week intervention phases separated by a one-week washout period to reduce carryover effects (**Figure 1A**). To assess the feasibility of down-PTAS while minimizing order bias, the SHAM and STIM condition weeks were randomized. Condition details were concealed from participants to prevent influencing natural sleep and from those involved in sleep stage scoring and data analysis to maintain objectivity. During the screening visit, after providing written informed consent, participants received the study devices and were familiarized with all study procedures, which they later conducted at home. To ensure inclusion criteria were fulfilled and no exclusion criteria were met, they completed a baseline assessment that included the Pittsburgh Sleep Quality Index (PSQI, [21]) to evaluate subjective sleep quality over the past month (inclusion threshold < 10), the Epworth Sleepiness Scale (ESS, [22]) to determine daytime sleepiness over the same period (inclusion threshold < 11), the 21-item Hamilton Depression Rating Scale (HDRS, [23]) for observer-rated depression symptoms over the past week (inclusion threshold < 8), and the Beck Depression Inventory-II (BDI-II, [24]) for self-rated depression symptoms over the past two weeks (inclusion threshold < 14).

**Figure 1.**
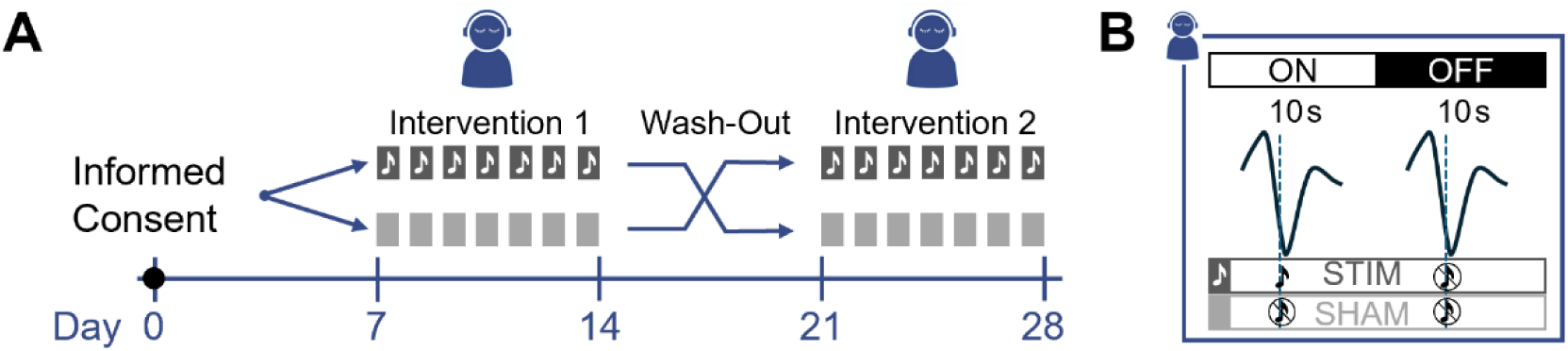
Study and stimulation protocols. **(A)** Schematic overview of a participant’s timeline. Written informed consent was obtained during the screening visit, followed by administration of the Karolinska Sleepiness Scale (KSS) three times daily – morning, afternoon, and evening – throughout the study period. Participants were randomly assigned to receive either the STIM condition first (dark grey with white note) followed by the SHAM condition (light grey), or vice versa. All interventions took place in a home setting. **(B)** Down-phase targeted auditory stimulation (down-PTAS) protocol. Stimulation was applied in a windowed manner during detected slow-wave sleep, alternating between 10-second ON windows, during which auditory stimulation was delivered just before the negative peak of slow waves, and 10-second OFF windows, where stimulation was disabled. In the SHAM condition, stimulation remained disabled during both ON and OFF windows.

Throughout the study, participants completed the Karolinska Sleepiness Scale (KSS, [25]) three times daily – in the morning, afternoon, and evening – via a smartphone questionnaire application based on the RADAR-Base platform [26]. During the intervention weeks, sleep was recorded using the MHSL-SB. Participants were guided through the device preparation in the evening and removal in the morning using checklists on the same smartphone application. Additionally, they were asked on two 5-point Likert scales about their perceived sleep quality (0 – very good to 4 – very bad) and sleep depth (0 – very superficial to 4 – very deep) of the preceding intervention night.

To address any questions or issues, a 24/7 helpline was provided, and participants were encouraged to contact the study team at any time. Remote data monitoring allowed for early intervention in case of technical difficulties.

### Home Sleep Recordings

Electroencephalography (EEG), electrooculography (EOG), and electromyography (EMG) data were collected using the MHSL-SB [27]. The frontal EEG channel (Fpz), two EOG channels, and two EMG channels were referenced to the right mastoid, while the left mastoid served as the ground electrode. Single-use, self-adhesive electrodes (BlueSensor N, Ambu A/S, DK) were employed for signal acquisition.

Data were recorded at a 250 Hz sampling rate with a 24-bit analog-to-digital converter and processed with a real-time anti-aliasing filter and a 50 Hz notch filter before being fed into the built-in stimulation algorithm. This algorithm performs a binary classification of EEG signals into NREM and non-NREM sleep based on spectral power thresholds in the low delta (2–4 Hz), high delta (3–5 Hz), and high beta (20–30 Hz) bands. Default power thresholds, derived from historical data of a similar age cohort, were applied consistently across stimulation nights.

Once ten minutes of stable NREM sleep were detected, stimulation was initiated. A phase-locked-loop algorithm [27] then identified slow waves in real time, triggering stimuli at a target phase of 225°, corresponding to the transition from the upstate to the downstate of the ongoing slow wave. The minimal inter-stimulus interval (ISI) was set to 0.5 s.

Stimulation followed a block design with alternating 10-second ON windows, during which stimulation was active, and 10-second OFF windows, where it was disabled (**Figure 1B**).

During SHAM nights, all stimuli were disabled. For STIM nights, stimuli in the OFF windows and for SHAM nights, stimuli in all windows were reconstructed offline using the algorithm’s power thresholds. Each stimulus consisted of a 50 ms burst of pink noise delivered through headphones embedded in a headband. The sound pressure level started at 50 dB and gradually increased until an arousal was detected, at which point it returned to the initial volume before increasing again. For safety considerations, the sound pressure level was set to never exceed 70 dB. Mean stimulation phase and standard deviation were extracted for each sleep recording using a toolbox for circular statistics [28] in MATLAB R2024b (The MathWorks Inc., 2024), and grand means and grand standard deviations were computed for SHAM and STIM recordings separately.

### Sleep Stage Scoring and Artifact Rejection

Sleep stage scoring was performed using Visbrain [29] (Python 3.8.10) following AASM guidelines [30]. Before visualization, 3rd-order Butterworth digital IIR filters were applied bidirectionally to prevent phase distortions. According to AASM recommendations [30], the Fpz signal underwent band-pass filtering between 0.5 Hz and 35 Hz, while the two EOG channels were filtered between 0.3 Hz and 35 Hz. Chin EMG signals were band-pass filtered between 10 Hz and 100 Hz and re-referenced bilaterally when both channels exhibited good signal quality upon visual inspection. After filtering, all signals were down sampled to 128 Hz, as required by the scoring software.

Vigilance states (W, N1, N2, N3, R) were visually scored in 30-second epochs by a sleep expert, with validation from a second expert. Both scorers were blinded to the experimental condition. Various sleep metrics were derived from the visual scoring.

Time in bed (TIB) was defined as the duration between the start and end of the recording or the removal of the headband, whichever occurred first. Total sleep time (TST) included all epochs within TIB that were not scored as W. Sleep efficiency was calculated as the percentage of TST relative to TIB. Percentages for each sleep stage (W, N1, N2, N3, REM) were also computed based on TIB. Wake after sleep onset (WASO) was defined as the total time spent in stage W after the first occurrence of any epoch not scored as W.

Following scoring, NREM sleep epochs (N1, N2, N3) underwent a semi-automatic artifact detection procedure adapted from Leach et al. [31]. The adaptation, required for using the tool with single-channel data, is detailed in [32].

### Electroencephalography Signal Preprocessing

The EEG signal was first processed with a low-pass filter, allowing frequencies up to 30 Hz (–6 dB attenuation at 39.86 Hz; filter order: 46, sampling rate: 250 Hz), followed by a high-pass filter with a pass band starting at 0.5 Hz (–6 dB attenuation at 0.37 Hz; filter order: 2986, sampling rate: 250 Hz), similar to a previous study from our laboratory [1]. Both filters were Kaiser window FIR filters with zero-phase shift to prevent phase distortions. Filtering and all subsequent signal processing steps were conducted in MATLAB R2024b (The MathWorks Inc., 2024) using EEGLAB [33].

### Slow-Wave Activity Analysis

Previous studies have shown that the effects of down-PTAS were most prominent in low-frequency ranges, either below 1 Hz [4] or up to 2 Hz [1,3]. Therefore, we calculated the average SWA in the low-frequency range (0.5–2 Hz, lSWA). For each 10-second ON and OFF window, power spectral density was computed using the MATLAB’s *pwelch*-function (Welch method) for 4-second Hanning windows with a 50% overlap, resulting in a 0.25 Hz frequency resolution, and lSWA was calculated as the power between 0.5-2 Hz.

First, we analyzed whole-night lSWA during NREM sleep (stages N2 and N3) in artifact-free windows combined. For a more focused evaluation of stimulation effects, next, we selected only ON windows that contained at least one stimulus, along with their corresponding consecutive OFF windows (stimulated ON-OFF window pairs), and analyzed them combined and for ON and OFF windows separately.

Following the approach of Leach et al. [1], we categorized the stimulated ON-OFF window pairs based on the number of stimuli within the ON window. Given the 10-second window length, 1–7 stimuli within ON windows were classified as *few*, while 8–14 stimuli were classified as *many*.

To complement the whole-night analyses, recordings were segmented into sleep cycles based on the method by Feinberg and Floyd [34], and lSWA was calculated for the first four NREM cycles. Sleep recordings with fewer than four completed cycles were excluded from this analysis. First, we analyzed all artifact-free windows during NREM sleep. As before, we then focused specifically on stimulated ON-OFF window pairs, conducting the analysis both combined and separately for ON and OFF windows.

### Time-Frequency Analysis

Time-frequency analyses were conducted using Morlet wavelets with 3 cycles at the lowest frequency (1 Hz) and up to 10 cycles at the highest frequency (25 Hz), with frequencies logarithmically spaced. This yielded phase and spectral power (event-related spectral perturbation, ERSP) values for 97 frequencies ranging from 1 to 25 Hz (in 0.25 Hz steps). First, the Fpz EEG data were synchronized to the stimulus onset of each stimulus in the ON window.

Then, the stimuli were divided into isolated (ISI ≥5 s) and consecutive (ISI ≤1 s) stimuli similar to Leach and co-authors [1]. For each stimulus, EEG data within a brief time-window (−3.4 to 5.604 s) were processed using a standard wavelet transformation routine [35]. To avoid edge artifacts, the first and last 3 seconds of data were excluded as buffer zones, resulting in phase and spectral power values for 751 time points between −0.4 and 2.6 s (at a sampling rate of 250 Hz). To reduce file size, these values were then down-sampled to 50 Hz. Finally, to correct for participant and frequency biases, spectral power values were normalized by dividing them by the average spectral power across the entire time window, calculated from the average of both conditions (SHAM and STIM), on a participant- and frequency-wise basis.

Subsequently, we repeated the same analysis time-locked to the first stimulus in the ON window, extracting a longer time window (−3.4 to 18.004 s). This yielded phase and spectral power values for 3’751 time points between −0.4 and 15 s. ON-OFF window pairs in which the first stimulus occurred in the second half of the ON window were excluded to ensure that time points between 10 and 15 s remained entirely within OFF windows.

For the same timeframes and stimuli, the average Fpz EEG was extracted for both conditions (SHAM and STIM), with their difference reflecting the auditory evoked response (AER).

### Arousal Detection

A robust automatic EEG arousal detection algorithm was applied, utilizing both EEG and EMG signals [36,37]. The Fpz EEG signal was fed into the algorithm after undergoing the preprocessing steps described above. EMG data underwent several stages of filtering: lowpass filtering with a −6 dB attenuation at 96.49 Hz (filter order: 130 at 250 Hz), notch filtering at 50 Hz (filter order: 184 at 250 Hz), and high-pass filtering with −6 dB attenuation at 14.01 Hz (filter order: 122 at 250 Hz). The EMG signals were then re-referenced bilaterally.

As with EEG preprocessing, Kaiser window FIR filters with zero-phase shift were applied. Arousals occurring during wakefulness (stage W) were excluded.

### Statistics

Due to multiple recordings per condition for each participant, linear mixed effects models were computed with the metric of interest as the dependent variable, condition (SHAM as reference vs. STIM) as the fixed effect, and random participant intercepts. When dividing the sleep recordings into sleep cycles, we treated sleep cycle as a factorial variable and included it as a fixed effect, added an interaction term with condition, and computed estimated marginal means (EMMs) for the STIM-SHAM contrast over all cycles and for each cycle separately. For the comparison of stimulation numbers and phase, sleep stage scoring metrics, arousals, subjective sleepiness (KSS), perceived sleep quality and depth, we report unadjusted *p*-values to achieve high sensitivity for detection of condition differences and potential side effects of down-PTAS. Unless otherwise specified, *p*-values (*p*) are FDR-corrected according to Benjamini and Hochberg [38], β-coefficients (*β*) or EMMs (*EMMs_STIM-SHAM_*) and Cohen’s d (*d*) are reported alongside, and data are presented as mean ± standard deviation (SD). Significant differences are reported for *p*<0.05. For exploratory analyses of this feasibility study, differences with *p*<0.1 are consistently reported and discussed as trends. In cases where there is divergence between unadjusted and FDR-corrected *p*-values, both are presented. All statistical analyses and visualizations, except for the time-frequency analysis, were performed using R Statistical Software version 4.4.1 [39] and the following packages: here, tidyverse, data.table, chron, writexl, ggpubr, rstatix, ggsignif, nlme, and emmeans.

The statistics for the time-frequency analysis was computed in MATLAB R2024b (The MathWorks Inc., 2024) using the lmeEEG package [40], which enables linear mixed effects models with mass univariate analyses, adapted for frequency-timepoint instead of channel-timepoint. Specifically, we first conducted linear mixed effects models with the ERSP values for each frequency-timepoint combination, again using condition as the fixed effect and random participant intercepts. In the next step, we extracted a “marginal” ERSP by removing random-effects contributions from the data. We then performed mass univariate linear regressions on the “marginal” ERSP data. Subsequently, we permuted the design matrix 2000 times to properly estimate critical statistics, using an α-level of 0.05. The permutation function was modified to generate 2000 overall unique permutations, yet allowing for duplicate assignments within participants. Finally, we applied threshold-free cluster enhancement (TFCE) to identify significant clusters.

## Results

### Data Quality of Unsupervised Home Sleep Recordings

Participant characteristics are shown in Table 1. In brief, the three women and two men were healthy young adults with good subjective sleep quality, normal daytime sleepiness, and neither observer- nor self-rated depressive symptoms. All participants completed the study consisting of 14 sleep recordings. A single adverse event was reported, involving mild, self-limiting common cold symptoms associated with a confirmed COVID-19 infection during the washout period after the SHAM condition. This adverse event did not prevent the participant from completing the protocol, and the study team was only retrospectively informed of this.

**Table 1.** Participant Characteristics.

|  |  | Mean±SD/Ratio |
| --- | --- | --- |
| Sex | [f/m] | 3/2 |
| Age | [years] | 23.8±0.8 |
| BMI | [kg/m<sup>2</sup>] | 22.6±2.8 |
| PSQI score |  | 4.2±1.6 |
| ESS score |  | 7.2±1.9 |
| HDRS score |  | 0.8±1.8 |
| BDI-II score |  | 4.6±4.0 |

*PSQI: Pittsburgh Sleep Quality Index; ESS: Epworth Sleepiness Scale; HDRS: Hamilton Depression Rating Scale; BDI-II: Beck Depression Inventory-II*

Of the 70 potential sleep recordings, 67 (96%) were available, with three missing due to technical issues. For three participants, the remote device parameter settings were only applied after the first sleep recording, resulting in the exclusion of these three recordings. Additionally, four recordings were excluded due to a technical issue causing a sudden termination of the recording, and two more were excluded due to signal loss exceeding 20% of the recording duration. This left 58 sleep recordings (87% of available data) for further analysis – 29 SHAM and 29 STIM nights.

For each sleep recording, the number of hypothetical (SHAM) and actual (STIM) stimuli during ON windows exceeded 700, with an average of 1759±522 stimuli (mean ± standard deviation) during SHAM and 1640±438 stimuli during STIM nights without evidence for a condition effect (*p_unadjusted_*=0.22). The grand mean stimulation phase was 238.7° for SHAM nights and 237.8° for STIM nights, with grand standard deviations of 59.8° and 69.1°, respectively. Similarly, there was no evidence for a difference between conditions (*p_unadjusted_*=0.12). The phase distribution of stimuli is visualized in **Supplementary** Figure 1.

### Sleep Macrostructure and Subjective Sleep Perception

Comprehensive sleep metrics are summarized in Table 2. There is no evidence of a difference between the two conditions in time spent in bed (TIB), sleep latency, sleep continuity measures (total sleep time, sleep efficiency, WASO), REM latency, and sleep stage distribution (Wakefulness, N1, N2, N3, REM). The same holds true for the number of arousals, regardless of whether considering α– and β-arousals separately or in combination.

**Table 2.** Sleep metrics.

| Sleep Metric | | SHAM | STIM | $p_{unadjusted}$ |
| --- | --- | --- | --- | --- |
| Time in bed (TIB) | [min] | 470±64 | 450±73 | 0.32 |
| Total sleep time | [min] | 446±60 | 425±65 | 0.24 |
| Sleep efficiency | [%TIB] | 94.9±2.7 | 94.7±2.9 | 0.31 |
| WASO | [%] | 1.7±1.4 | 2.0±1.3 | 0.33 |
| Sleep latency | [min] | 16±11 | 16±11 | 0.84 |
| REM latency | [min] | 98±33 | 98±34 | 0.70 |
| N1 sleep | [%TIB] | 3.4±1.9 | 3.5±1.6 | 0.42 |
| N2 sleep | [%TIB] | 44.1±5.7 | 44.1±7.0 | 0.75 |
| N3 sleep | [%TIB] | 22.6±4.8 | 23.0±7.1 | 0.91 |
| REM sleep | [%TIB] | 24.1±4.6 | 23.4±4.1 | 0.50 |
| Wakefulness | [%TIB] | 5.1±2.7 | 5.3±2.9 | 0.31 |
| Arousals combined |  | 100.8±35.9 | 96.8±38.6 | 0.76 |
| β-Arousals | | 89.8±33.2 | 87.6±36.3 | 0.94 |
| α-Arousals | | 11.0±6.9 | 9.2±6.4 | 0.17 |
| Morning KSS |  | 2.9±1.4 | 3.6±1.6 | 0.09 |
| Afternoon KSS |  | 2.7±1.9 | 2.8±2.1 | 0.60 |
| Evening KSS |  | 5.9±1.6 | 5.6±1.6 | 0.55 |
| $\Delta KSS_{Morning - Evening}$ | | -2.9±1.8 | -2.0±1.8 | 0.05 |
| Sleep Quality |  | 2.3±0.8 | 2.4±1.1 | 0.49 |
| Sleep Depth |  | 2.6±0.8 | 2.5±0.7 | 0.84 |

WASO: Wake after sleep onset, calculated as percentage of time spent awake after the first occurrence of any epoch not scored as wakefulness; KSS: Karolinska Sleepiness Scale; Perceived sleep quality and depth from 5-point Likert scales (0 – very good quality; 0 – very superficial sleep depth); numbers represent means ± standard deviations; for higher sensitivity to detect potential differences, unadjusted $p$ -values ( $p_{unadjusted}$ ) derived from linear mixed effects models with random participant intercepts are reported.

Regarding subjective sleepiness measured by the KSS, unadjusted *p*-values indicate a trend toward higher morning values and a lower overnight decrease (ΔKSS_Morning-Evening_) after STIM compared to SHAM nights, without evidence for a condition difference in the afternoon and evening and for any timepoint or overnight decrease after FDR-correction (*p_FDR_* > 0.15). A closer single-day comparison, including the baseline (BL) week before any sleep intervention, is shown in **Supplementary** Figure 2. Marked inter– and intra-individual variability is evident across all three conditions, with no signs of worsening over the intervention weeks that would suggest an accumulation of relevant sleepiness. As for perceived sleep quality and depth, there was no evidence of a difference between STIM and SHAM nights.

### Electrophysiological Stimulus Response

When examining whole-night lSWA (0.5-2 Hz, see Methods) in artifact-free NREM sleep (stages N2 and N3), there is no evidence of a difference between conditions (*β* = −1.0, *p* = 0.93, *d* = 0.02; **Figure 2A**, Whole-Night). For roughly half of the ON-OFF window pairs in artifact-free NREM sleep (50.5%), there are no stimuli delivered during the ON window (data not shown). To evaluate stimulation effects more closely, we therefore focused specifically on ON-OFF window pairs containing at least one stimulus in the ON window (**Figure 2A**, ON-OFF St.). For these stimulated ON-OFF window pairs, lSWA shows a trend towards lower values during STIM as compared to SHAM nights (*β* = −51.6, *p* = 0.052, *d* = 0.61), with a significant difference before FDR correction (*p* = 0.026). Further separation into ON and OFF windows reveals no difference between conditions in ON windows (*β* = 17.4, *p* = 0.66, *d* = −0.17, **Figure 2A**, ON St.), but a significantly lower lSWA in OFF windows during STIM nights (*β* = −121.0, *p* < 0.0001, *d* = 1.37; **Figure 2A**, OFF St.).

**Figure 2.**
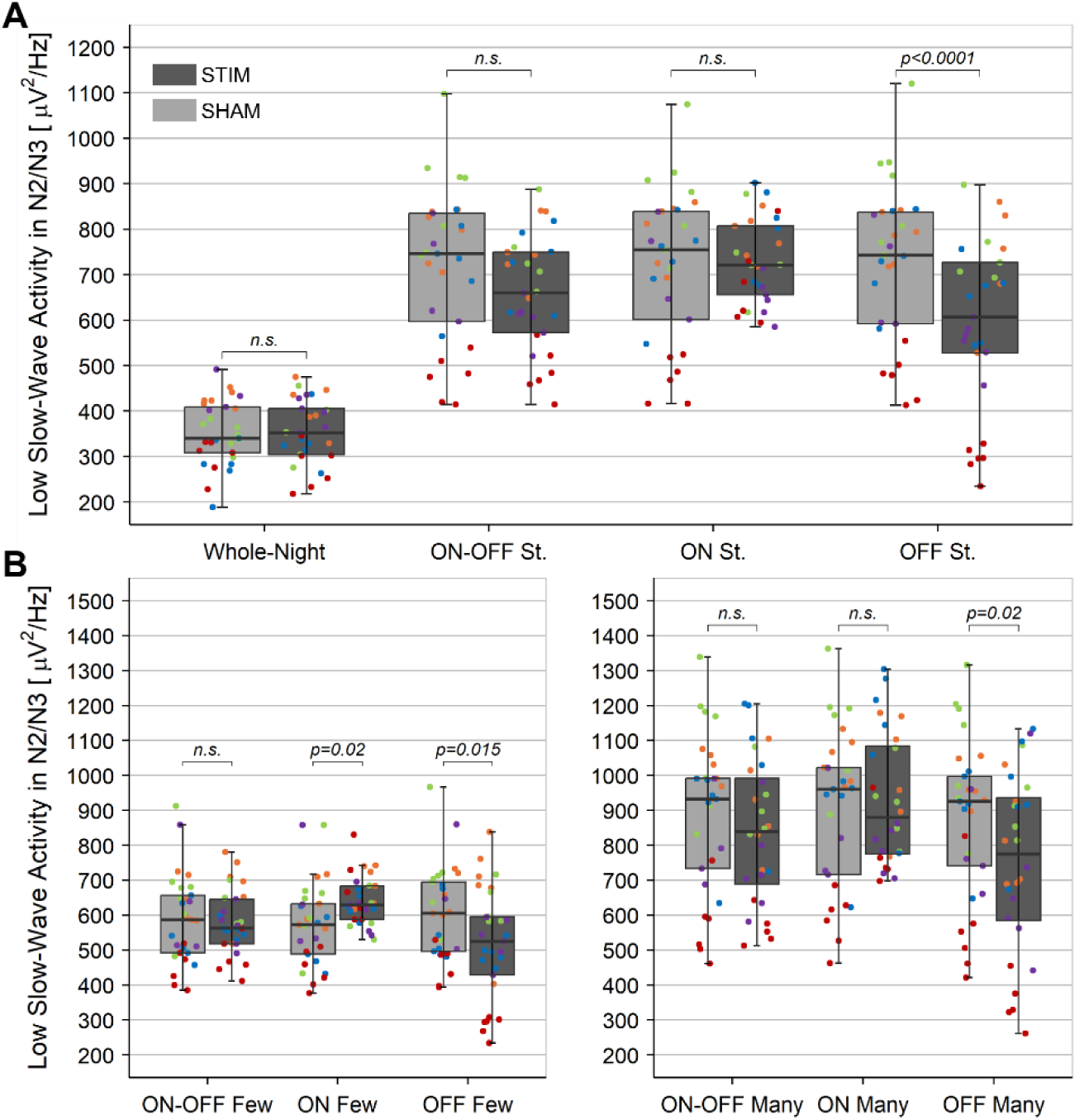
Low Slow-Wave Activity (lSWA, power between 0.5–2 Hz) during NREM sleep (stages N2 and N3). **(A)** Results are shown for whole-night artifact-free windows (Whole-Night), ON-OFF window pairs with at least one stimulus in the ON window (ON-OFF Stimulated), and the respective ON (ON Stimulated) and OFF (OFF Stimulated) windows separately. **(B)** Stimulated ON-OFF window pairs are further categorized into those with few (1–7) and many (8–14) stimuli in the ON window. Individual participant data (n=5) are color-coded, with each dot representing the mean of one sleep recording (n=29 SHAM, n=29 STIM nights). SHAM nights are depicted in light grey, STIM nights in dark grey boxes. The horizontal line marks the median, box hinges represent the first and third quartiles, whiskers extend to the largest value within 1.5 times the interquartile range, and outliers are individually plotted. Significant FDR-corrected p-values from linear mixed-effects models are shown above; n.s. indicates p-values >0.05.

Due to the differential stimulus responses depending on the number of stimuli delivered during the ON window observed by Leach et al. [1], we examined ON-OFF window pairs with few (1–7) and many (8–14) stimuli separately (**Figure 2B**). For ON windows with few stimuli, lSWA increases significantly (*β* = 68.7, *p* = 0.02, *d* = −0.73), followed by a decrease in subsequent OFF windows (*β* = −79.2, *p* = 0.015, *d* = 0.82), with no overall change when combining ON and OFF windows (*β* = −5.3, *p* = 0.90, *d* = 0.06). In contrast, ON windows with many stimuli show no evidence of a condition effect (*β* = 34.4, *p* = 0.57, *d* = −0.23), but OFF windows again exhibit a significant lSWA reduction in STIM nights (*β* = −112.2, *p* = 0.02, *d* = 0.74). As before, no difference is observed when combining ON and OFF windows (*β* = −38.8, *p* = 0.50, *d* = 0.28).

As illustrated in **Figure 2B**, lSWA appears higher in window pairs with many stimuli compared to those with few stimuli, representative for deeper sleep in windows with many stimuli. Since sleep depth typically declines across the night, and also due to the potential for device misplacement over the course of the night, we segmented the nights into sleep cycles to assess whether stimulation effects varied throughout the night, again beginning with all artifact-free windows in N2 and N3 sleep and moving on to stimulated ON-OFF windows combined as pairs and separate for ON and OFF windows (**Figure 3**).

**Figure 3.**
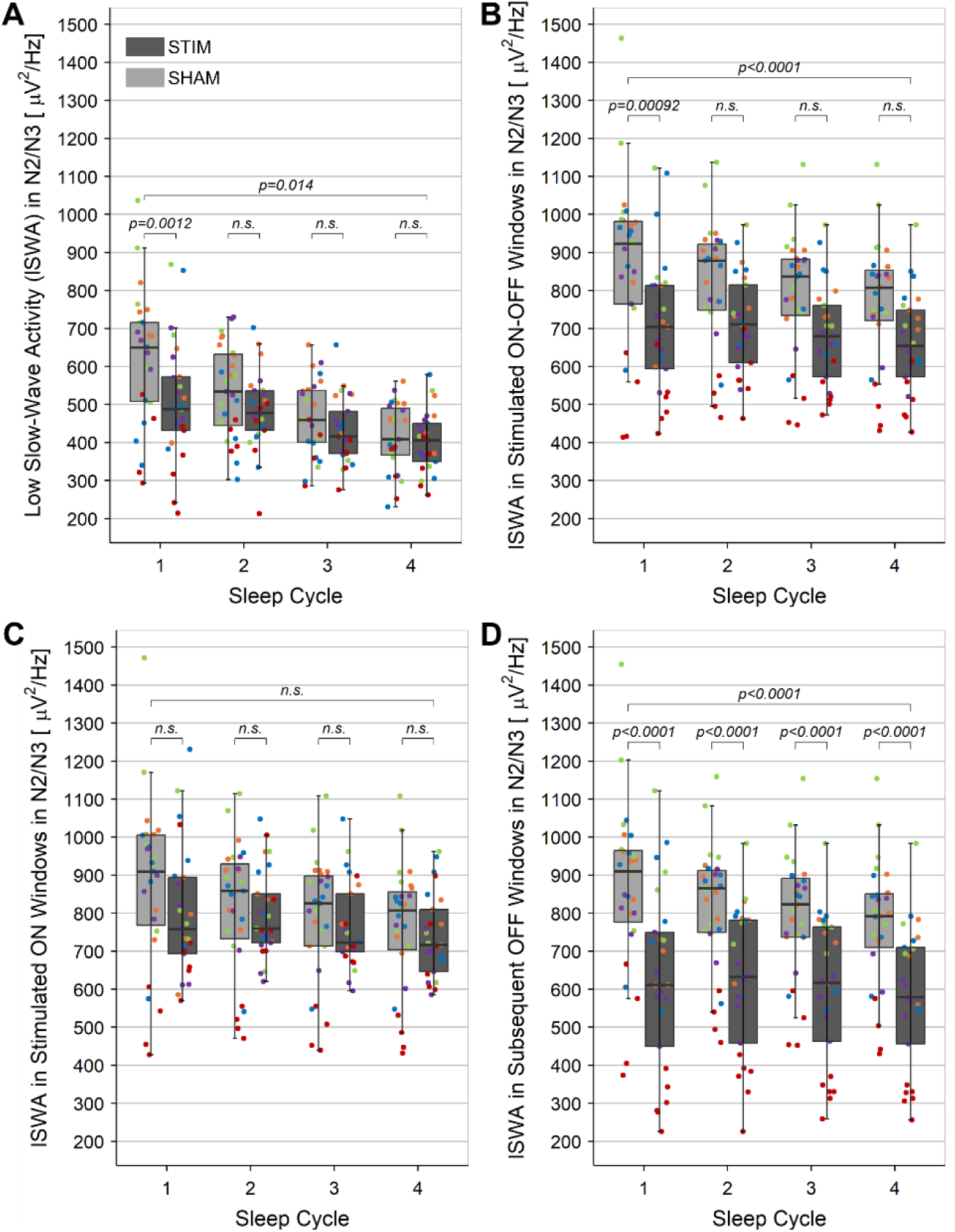
Low Slow-Wave Activity (lSWA, power between 0.5–2 Hz) during NREM sleep (stages N2 and N3), segmented into sleep cycles based on Feinberg and Floyd [34]. The first four sleep cycles are shown for **(A)** all artifact-free windows, **(B)** ON-OFF window pairs with at least one stimulus in the ON window, **(C)** stimulated ON windows, and **(D)** subsequent OFF windows. Data presentation, color-coding, and significance indications follow the conventions in **Figure 2**. Six sleep recordings of four participants lacked four complete sleep cycles and were therefore excluded from the analysis, resulting in a final sample of n = 52 analyzed recordings.

Accounting for sleep cycles, lSWA is significantly lower in STIM compared to SHAM nights when including all artifact-free windows in NREM sleep (*EMMs_STIM-SHAM_* = −39.0, *p* = 0.01, *d* = 0.39; **Figure 3A**) and stimulated ON-OFF window pairs only (*EMMs_STIM-SHAM_* = −77.2, *p* < 0.0001, *d* = 0.68; **Figure 3B**). Post hoc tests indicate that this difference is driven by the first sleep cycle, where lSWA is significantly lower during STIM nights in both all artifactfree windows (*EMMs_STIM-SHAM_* = −98.6, *p* = 0.001, *d* = 0.98; **Figure 3A**) and stimulated ON-OFF window pairs (*EMMs_STIM-SHAM_* = −116.2, *p* = 0.001, *d* = 1.16; **Figure 3B**).

For later sleep cycles, all artifact-free windows (**Figure 3A**) show no evidence of a condition difference (|*EMMs_STIM-SHAM_* | < −40, *p* > 0.15, |*d*| < 0.40). In stimulated ON-OFF window pairs (**Figure 3B**), trends toward lower lSWA in STIM nights emerge in cycles 2 (*EMMs_STIM-SHAM_* = −61.8, *p* = 0.09, *d* = 0.62), 3 (*EMMs_STIM-SHAM_* = −64.2, *p* = 0.08, *d* = 0.64) and 4 (*EMMs_STIM-SHAM_* = −66.8, *p* = 0.07, *d* = 0.67), with significance before FDR correction in cycles 3 and 4 (*p* = 0.044 and *p* = 0.036, respectively).

Further subdivision into ON (**Figure 3C**) and OFF (**Figure 3D**) windows reveal no evidence for a difference in ON windows, neither over all four nor for individual cycles (|*EMMs_STIM-SHAM_* | < −50, *p* > 0.30, |*d*| < 0.50), but a significant reduction of lSWA in OFF windows during STIM nights as compared to SHAM nights across the four cycles together and individually (|*EMMs_STIM-SHAM_* | > −135, *p* < 0.0001, |*d*| > 1.3).

To evaluate the feasibility of replicating the stimulation response observed in highly controlled sleep laboratory settings [1] in an unsupervised home environment, a time-frequency representation and an auditory evoked response (AER) time-locked to stimulus onset are presented in **Figure 4**. Consistent with Leach et al. [1], averaging the Fpz EEG time-locked to stimulus onset revealed a prominent AER (black line), characterized by the classical sleep AER components P200, N550, and P900 [41,42]. As observed in this previous research [1], the N550 component is most pronounced in isolated stimuli with ISI ≥5 s (**Figure 4B**) and first stimuli with ISI ≥10 s (**Figure 4D**). This results in a more prominent and widespread increase in low slow-wave activity (lSWA) around 0.5 s and an increase in sigma activity (12– 16 Hz) around 0.9–1.5 s after stimulus onset. While these differences between hypothetical (SHAM) and actual (STIM) stimuli are also present, albeit to a lesser extent, when averaging across all stimuli (**Figure 4A**), they are notably attenuated for consecutive stimuli with ISI ≤1 s (**Figure 4C**). The longer timeframe in **Figure 4D** highlights the first significant cluster with reduced lSWA appearing early in the OFF window (∼10.2 s), with a more pronounced effect between 13–15 s after the onset of the first stimulus.

**Figure 4.**
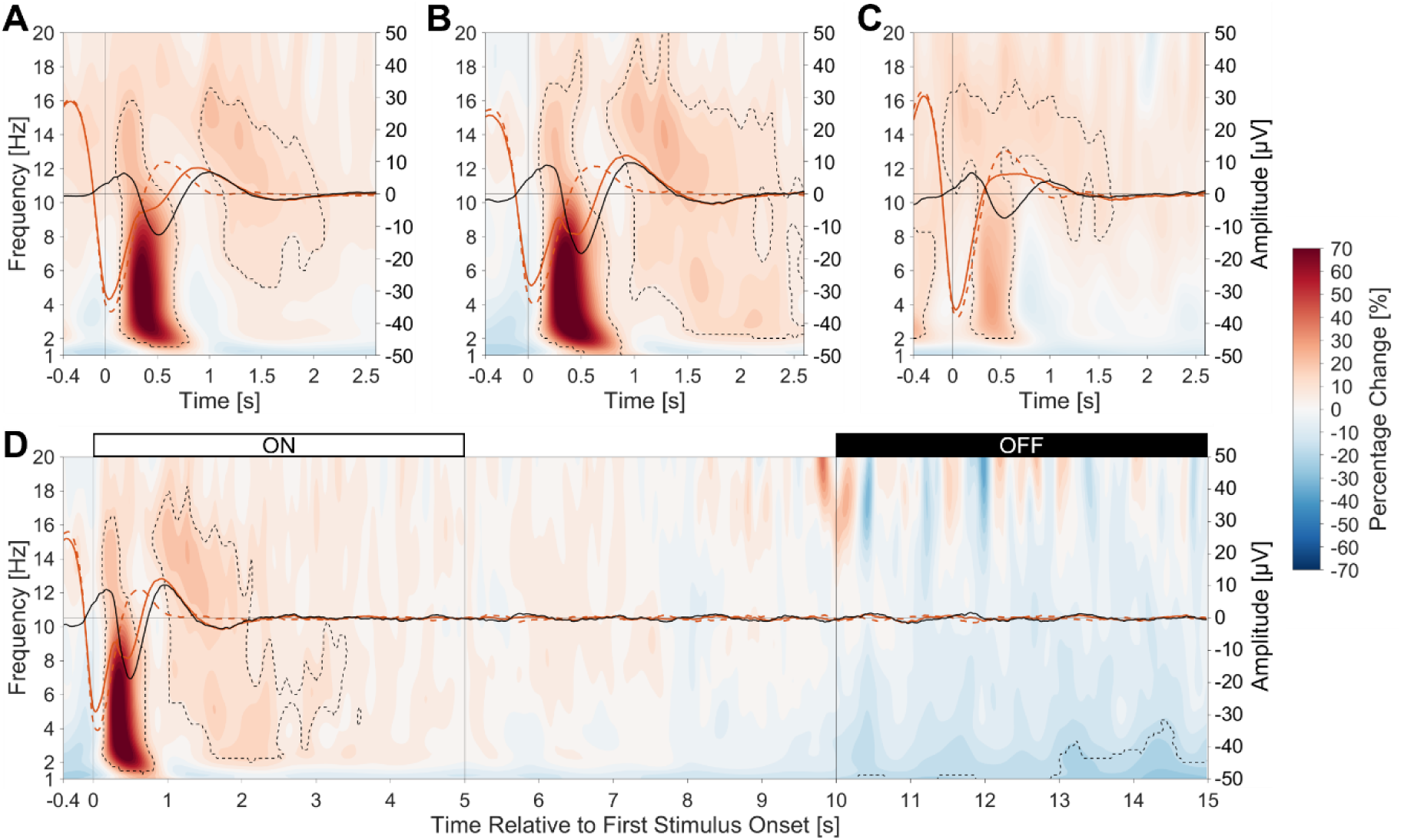
Averaged Fpz EEG responses to stimuli (right y-axis), time-locked to actual (STIM, solid red line) and hypothetical (SHAM, dashed red line) stimuli, and auditory evoked response (difference between the two, black line). Event-related spectral perturbation (ERSP) is shown in the background (left y-axis; normalized per participant and frequency using the trial average across the entire time window). Dashed contour lines indicate significant differences between conditions (linear mixed-effects models with mass univariate analyses and threshold-free cluster enhancement; see Methods). The top row presents data time-locked to all stimuli **(A)**, isolated stimuli with an interstimulus interval (ISI) ≥5 s **(B)**, and consecutive stimuli with an ISI ≤1 s **(C)**. Data timelocked to the first stimulus in the ON window is shown in **(D)**, restricted to cases where the first stimulus occurs within the first 5 s of the ON window, ensuring that the 0–5 s period falls entirely within the ON window and the 10–15 s period lies entirely within the OFF window.

## Discussion

Our findings demonstrate the successful application of down-PTAS in a home setting, highlighting its technical feasibility, favorable tolerability over seven consecutive nights, and robust delivery of auditory stimuli despite uncontrolled conditions. The high quality of the data in combination with the substantial number of auditory stimuli presented within the expected range of the targeted phase further validate the technical implementation of down-PTAS in an unsupervised home environment. Notably, nearly 90% of the available sleep recordings in our sample were of sufficient quality to permit manual sleep stage scoring and quantitative EEG analysis. All participants adhered to the study protocol, each completing 14 sleep recordings without reporting adverse events likely linked to the use of the MHSL-SB. No significant differences were observed between conditions regarding perceived sleep quality, sleep depth, and subjective sleepiness (KSS). This clearly indicates that seven consecutive nights of down-PTAS or SHAM-PTAS were well tolerated by healthy young adults.

In evaluating the feasibility of the selected device settings in generating comparable electrophysiological responses to those observed in controlled laboratory settings [1,3,4], our results revealed that while stimulation did not alter sleep architecture in our sample, it successfully modulated lSWA, reducing it particularly at the beginning of the night when sleep pressure and lSWA were still high, and specifically during OFF windows throughout the night. The absence of a whole-night lSWA reduction following down-PTAS aligns well with a number of previous studies [1,3,4]. For instance, Ngo *et al.* observed a rapid recovery of EEG power below 1 Hz after stimulation termination, which the authors attributed to the intrinsic drive of cortical networks to oscillate at their natural frequency below 1 Hz [4]. By contrast, Fattinger *et al.* reported continuous down-PTAS-induced reductions in lSWA [18]. These differences may be explained by variations in the stimulation protocol settings. While continuous down-PTAS may suppress lSWA, the windowed approach [1,3] and the 2.5 s discontinuation after two stimuli [4] likely result in more isolated stimuli associated with a K-complex-like response seen by Leach et al. [1], particularly at the beginning of ON windows or after breaks in stimulation.

The low-frequency component of this K-complex-like response may obscure a potential reduction in lSWA during acute stimulation in ON windows. This is further supported by our findings showing differential effects of stimulation during ON windows, depending on the number of stimuli delivered and the analysis of data across different sleep cycles. These observations suggest that setting the device to stimulate only the beginning of the night might be a tempting strategy. The choice of appropriate stimulation settings, however, should be guided by the intended use case. Given the evidence for a rebound after stimulation cessation [2,4], stopping the stimulation after the first sleep cycle could lead to an increase in lSWA in the subsequent cycles. This suggests that for clinical applications such as the treatment of depression – where the aim is to reduce lSWA to mimic aspects of sleep deprivation – continuous stimulation throughout the night may be necessary to maintain consistent effects across all cycles and avoid a rebound effect that could potentially prevent therapeutic benefits.

The time-frequency representation time-locked to stimuli closely mirrors findings by Leach et al. [1], suggesting that isolated stimuli likely induce a K-complex-like response, posing the question whether this would limit the therapeutic potential in depression. K-complexes are thought to have a dual role, either promoting sleep or facilitating arousal, depending on brain regions and the presence of a microarousal [43]. Rather than directly contributing to restorative sleep processes [44], K-complexes have been linked to various functions, including sleep protection in response to non-threatening stimuli [45,46], certain stages of memory replay and consolidation [47–49], and pathological conditions such as epilepsy [50], Alzheimer’s disease [51], and schizophrenia [52]. In the context of a potential use of the device with current settings for down-PTAS in depression, we assume that the observed K-complex-like responses may not diminish the intended effects but could complicate data interpretation by masking reductions in lSWA.

The observed significant reduction in lSWA following down-PTAS when analyzing sleep cycles, but not when averaging across the entire night, may be related to the unsupervised home setting. Sleep cycles potentially offer a more consistent reference for comparing nights when bedtimes and sleep opportunities were not strictly controlled. By examining specific sleep cycles across conditions, variability may be reduced, improving the statistical power to detect stimulation effects.

Several limitations should be considered when interpreting these findings. Despite collecting up to 14 sleep recordings per participant, this feasibility study involved only five healthy young adults with good sleep quality, limiting generalizability and warranting further research in a larger sample. From an ethical perspective, however, feasibility studies should minimize participant numbers. The home setting provides a significant advantage for longterm recordings and real-world data collection, which is valuable for potential clinical applications. This advantage, though, comes at the cost of increased variability compared to controlled laboratory settings. Nonetheless, the fact that our results closely resemble those observed in laboratory studies [7,9] suggests successful stimulation, and the higher number of recordings per condition may help balance the increased variability. Another limitation is the use of single-channel EEG recording, which prevents the assessment of topographical changes. Future studies should address this limitation by incorporating high-density EEG recordings in the laboratory alongside home studies.

## Conclusion

Recording sleep and performing down-PTAS over seven consecutive nights in an unsupervised home setting using the MHSL-SB is feasible, providing the foundation for future clinical trials. The therapeutic potential of this approach in the treatment of depression, however, can only be properly assessed in a patient population, as the effects of down-PTAS may differ significantly in individuals with clinical conditions. Testing in such a population is crucial to determine whether the promising results in terms of technical feasibility, tolerability, and EEG response to down-PTAS observed in healthy participants translate into meaningful therapeutic benefits for patients.

## Conflict of Interest

Reto Huber is co-founder and shareholder of Tosoo AG, a company focused on developing wearables for sleep electrophysiology monitoring and stimulation. Giulia Da Poian is board president of Tosoo AG. Hans-Peter Landolt serves as a compensated member of the Science Advisory Board for Heel Biologische Arzneimittel GmbH. Tosoo AG and Heel Biologische Arzneimittel GmbH had no involvement in any aspect of the work presented in this manuscript. The remaining authors report no competing interests.

## Data Availability

The data is securely stored on servers located at ETH Zurich. Ethics approval and guidelines restrict the online sharing of raw data. Access to the data, however, will be possible following a data sharing agreement combined with ethics approval on an individual user basis. The authors are happy to help researchers to access to this dataset.

## Funding Sources

This study was financially supported by Swiss National Science Foundation (SNSF) under grant numbers PZ00P2_193291 and 323530_207034 and supported by the Flagship Sleep-Loop grant of “Hochschulmedizin Zürich”, Switzerland.

## CRediT Author Statement

Corinne Eicher: Conceptualization, Methodology, Formal Analysis, Investigation, Writing – Original Draft, Visualization, Funding Acquisition. Cristina Gallego Vázquez: Data Curation, Writing – Review & Editing. Cinzia Schmid: Investigation, Writing – Review & Editing. Golo Kronenberg: Writing – Review & Editing. Hans-Peter Landolt: Conceptualization, Writing – Review & Editing. Erich Seifritz: Conceptualization, Writing – Review & Editing, Funding Acquisition. Giulia Da Poian: Conceptualization, Methodology, Data Curation, Writing – Review & Editing, Supervision, Project Administration, Funding Acquisition. Reto Huber: Conceptualization, Methodology, Writing – Review & Editing, Supervision, Funding Acquisition.

## Acknowledgments

We would like to express our gratitude to all our participants for their time, reliability, and valuable data contributions. We also thank Julian Amacker for his advice on implementing linear mixed effects models with mass univariate analyses using an adapted permutation function, and to all members of the Huber Sleep Lab for their ongoing support throughout the project.

