## SupplementaryMaterial for "Application of Down-Phase Targeted Auditory Stimulation During Sleep in a Home Setting: A Feasibility Study Across Seven Consecutive Nights"

### Supplementary Material

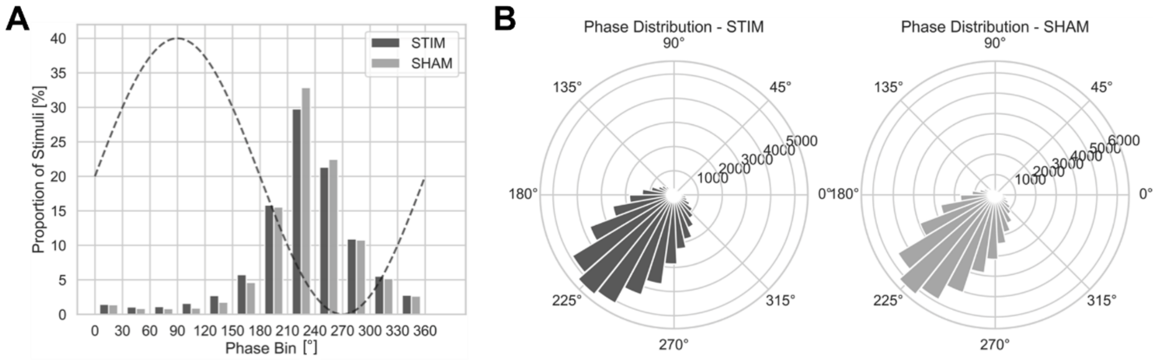


***Supplementary Figure 1.*** *Distribution of stimuli pooled across participants and nights within the same condition.* ***(A)*** *Bar plot showing the proportion of stimuli in each 30° phase bin, with a dashed sinusoidal curve representing the oscillation phase.* ***(B)*** *Polar histograms illustrating the phase distribution of all stimuli. Conditions are color-coded, with dark grey bars representing STIM and light grey bars representing SHAM.*

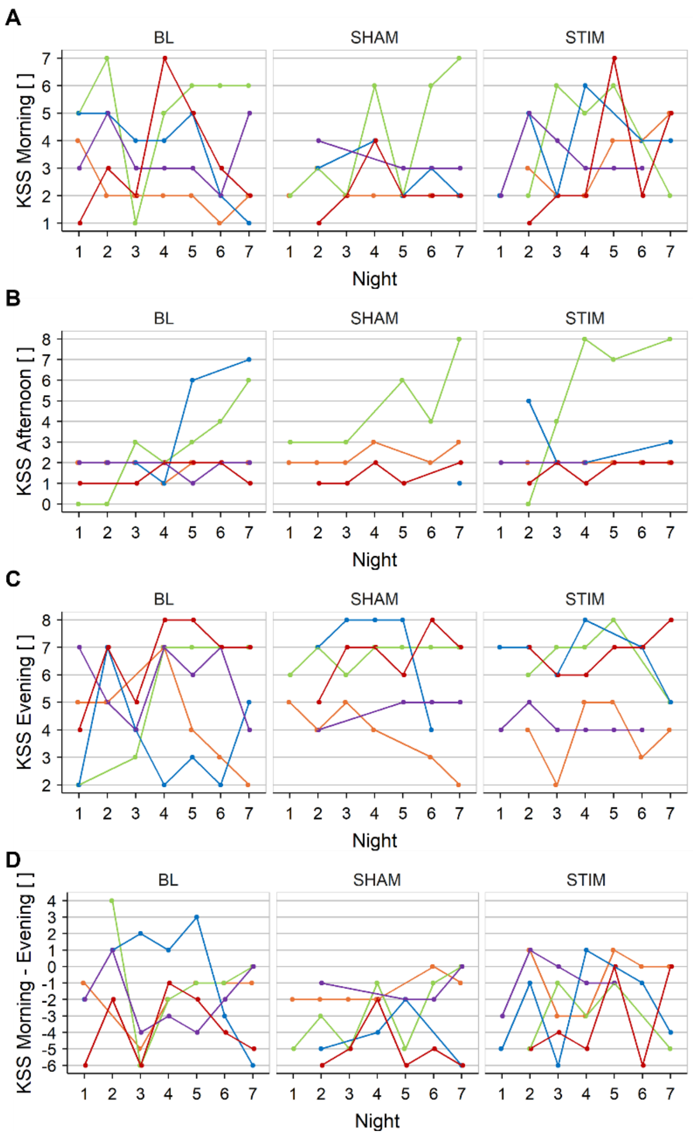


***Supplementary Figure 2.*** *Subjective sleepiness measured by the Karolinska Sleepiness Scale (KSS) across baseline (BL) and intervention (SHAM, STIM) weeks. The figure presents responses for morning **(A)**, afternoon **(B)**, and evening **(C)** assessments, as well as overnight changes **(D)**. Individual participants are color-coded and connected. The baseline week corresponds to the week immediately following the screening visit, before any sleep intervention.*
